# High-throughput single-cell DNA sequencing of AML tumors with droplet microfluidics

**DOI:** 10.1101/203158

**Authors:** Maurizio Pellegrino, Adam Sciambi, Sebastian Treusch, Robert Durruthy-Durruthy, Kaustubh Gokhale, Jose Jacob, Tina X. Chen, William Oldham, Jairo Matthews, Hagop Kantarjian, P. Andrew Futreal, Keyur Patel, Keith W. Jones, Koichi Takahashi, Dennis J. Eastburn

## Abstract

To enable the characterization of genetic heterogeneity in tumor cell populations, we developed a novel microfluidic approach that barcodes amplified genomic DNA from thousands of individual cancer cells confined to droplets. The barcodes are then used to reassemble the genetic profiles of cells from next generation sequencing data. Using this approach, we sequenced longitudinally collected AML tumor populations from two patients and genotyped up to 62 disease relevant loci across more than 16,000 individual cells. Targeted single-cell sequencing was able to sensitively identify tumor cells during complete remission and uncovered complex clonal evolution within AML tumors that was not observable with bulk sequencing. We anticipate that this approach will make feasible the routine analysis of heterogeneity in AML leading to improved stratification and therapy selection for the disease.

## Introduction

Current tumor sequencing paradigms are inadequate to fully characterize many instances of AML (acute myeloid leukemia) (*1, 2*). A major challenge has been the unambiguous identification of potentially rare and genetically heterogeneous neoplastic cell populations, with subclones capable of critically impacting tumor evolution and the acquisition of therapeutic resistance (*3–5*). Standard bulk population sequencing is often unable to identify rare alleles or definitively determine whether mutations co-occur within the same cell. Single-cell sequencing has the potential to address these key issues and transform our ability to accurately characterize clonal heterogeneity in AML; however, previous single-cell studies examining genetic variation in AML have relied upon laborious, expensive and low-throughput technologies that are not readily scalable for routine analysis of the disease.

An established approach for high-throughput and scalable single-cell sequencing uses molecular barcodes to tag the nucleic acids of individual cells confined to emulsion droplets (*6–9*). Although it is now feasible to perform single-cell RNA-Seq on thousands of cells using this type of approach, high-throughput single-cell DNA genotyping using droplet microfluidics has not been demonstrated on eukaryotic cells. This is primarily due to the challenges associated with efficiently lysing cells, freeing genomic DNA from chromatin and enabling efficient amplification in the presence of high concentrations of crude lysate (*10, 11*).

In this report, we present a microfluidic approach, relying on cell-identifying molecular barcodes, that overcomes existing barriers to high-throughput single-cell DNA sequencing. We focus our single-cell sequencing analysis on 62 genomic loci implicated in the acquisition or progression of AML. As a demonstration of the technology we characterize longitudinal samples from multiple AML patients and uncover features of clonal architecture that are not available from bulk sequencing data. This rapid, cost effective and scalable approach promises to make routine analysis of genetic variation in tumors a reality.

## Results & Discussion

### Droplet workflow for genomic DNA amplification and barcoding

To enable the characterization of genetic diversity within cancer cell populations, we developed a novel two-step microfluidic droplet workflow that enables efficient and massively-parallel single-cell PCR-based barcoding (Fig. 1a,b). The microfluidic workflow first encapsulates individual cells in droplets, lyses the cells and prepares the lysate for genomic DNA amplification using proteases. Following this lysate preparation step, the proteases are inactivated via heat denaturation and droplets containing the genomes of individual cells are paired with molecular barcodes and PCR amplification reagents. To demonstrate the advantage of the protease in the two-step workflow, we performed droplet-based single-cell TaqMan PCR reactions targeting the SRY locus on the Y chromosome, present as a single copy in a karyotypically normal cell (Fig. 2a). We carried out PACS (PCR-activated cell sorting) on calcein violet stained DU145 prostate cancer cells encapsulated and lysed with or without the addition of a protease (*12, 13*). In the absence of protease during cell lysis, only 5.2% of detected DU145 cells were positive for TaqMan fluorescence. The inclusion of the protease resulted in a dramatically improved SRY locus detection rate of 97.9%.

**Figure 1.**
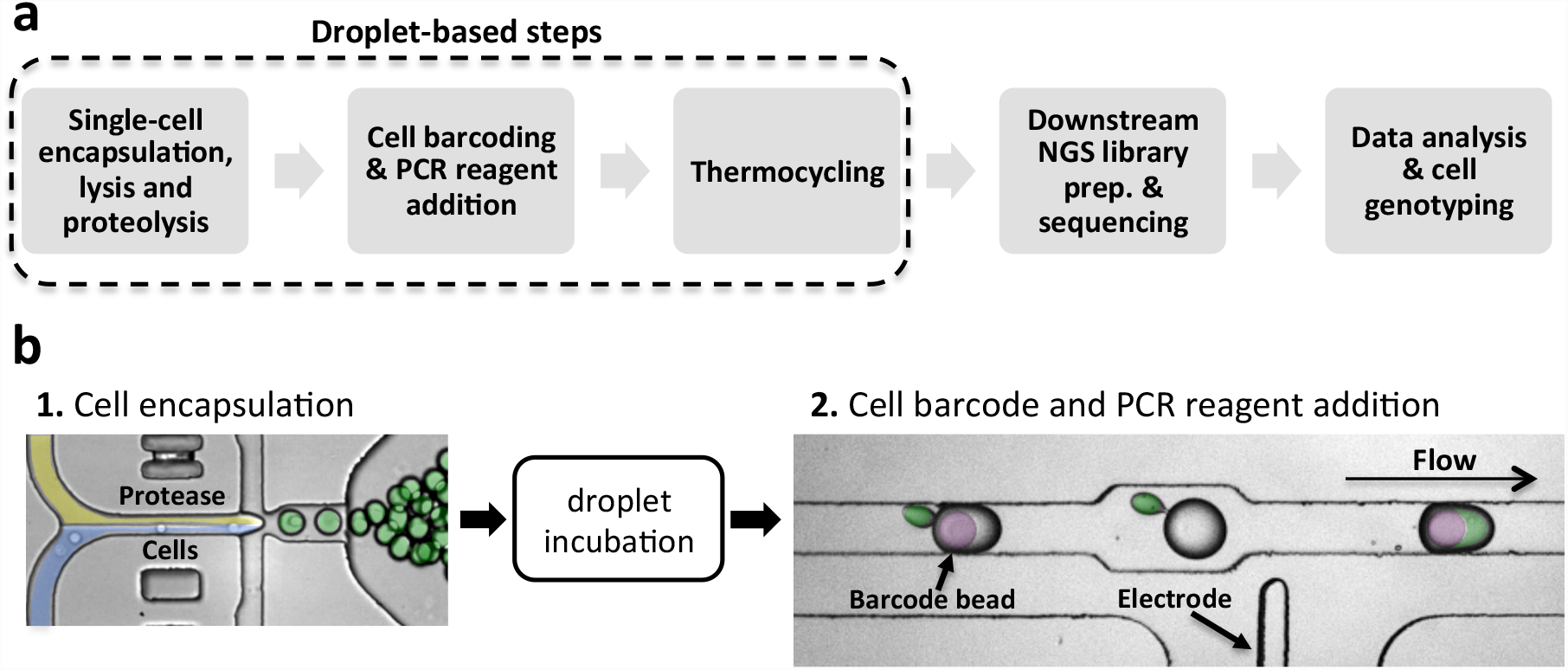
Protease based droplet workflow for single-cell genomic DNA amplification and barcoding. (a) Overview of the steps in the workflow. (b) Microfluidic devices to perform the two-step droplet workflow. Cells (pseudo colored in blue) are first encapsulated with lysis buffer containing protease (yellow) and incubated to promote proteolysis (green droplets). Protease activity is then thermally inactivated and the droplets containing the cell lysate are paired and merged with droplets containing PCR reagents and molecular barcode-carrying hydrogel beads (pseudo colored in purple).

**Figure 2.**
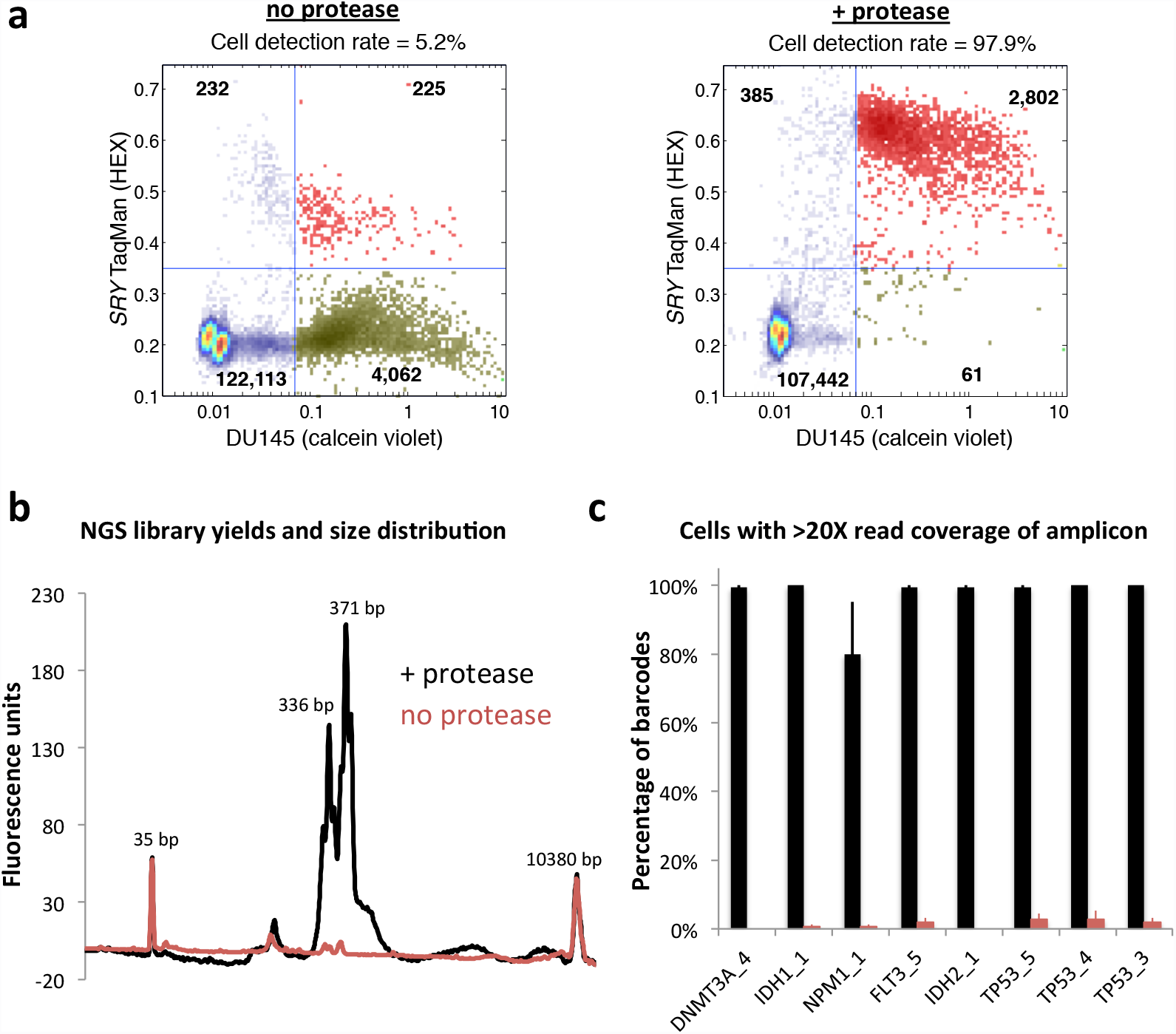
Protease-based workflows provide improved genomic DNA amplification. (a) When protease enzyme is left out of the workflow for single-cell gDNA PCR in droplets, only ~5% of DU145 cells (viability stained on the x-axis) are positive for SRY TaqMan reaction fluorescence (y-axis). Using protease during cell lysis improves the DU145 cell detection rate to ~98% (red points in upper right quadrant). Points in the plot represent droplets. (b) Bioanalyzer traces of sequencing libraries prepared from cells processed through the workflow with (black trace) or without (red trace) the use of protease indicates that PCR amplification in droplets is improved with proteolysis. The two-step workflow with protease enables better sequencing coverage depth per cell across the 8 amplified target loci listed on the x-axis (c).

We next sought to determine if the two-step workflow was also required for single-cell barcoding of amplicons targeting 8 genomic loci located in *TP53*, *DNMT3A*, *IDH1*, *IDH2*, *FLT3* and *NPM1*. To do this we synthesized hydrogel beads with oligonucleotides containing both cell identifying barcodes and different gene specific primer sequences (*6*). These barcoded beads were microfluidically combined with droplets containing cell lysate generated with or without the protease reagent (Fig 1b). Prior to PCR amplification, the oligonucleotides are photo-released from the hydrogel supports with UV exposure. Consistent with our earlier single-cell TaqMan reaction observations, amplification of the targeted genomic loci was substantially improved by use of a protease during cell lysis. Although similar numbers of input cells were used for both conditions, the use of protease enabled greater sequencing library DNA yields as assessed by a Bioanalyzer (Fig. 2b). Moreover, following sequencing, the average read coverage depth for the 8 targets from each cell was considerably higher when protease was used in the workflow (Fig. 2c). This data demonstrates the advantage of the two-step workflow for efficient amplification across different genomic loci for targeted single-cell genomic sequencing with molecular barcodes.

### Targeted sequencing of AML tumor samples

Having developed the core capability to perform targeted single-cell DNA sequencing, we next sought to apply the technology to the study of clonal heterogeneity in the context of normal karyotype AML. To provide variant allele information at clinically meaningful loci, we developed a 62 amplicon targeted panel that covers many of the 23 most commonly mutated genes associated with AML progression (Supplementary Table 1) (*14, 15*). Following optimization for uniformity of amplification across the targeted loci (Supplementary Figure 1), this panel was then used for single-cell targeted sequencing on AML patient bone marrow aspirates collected longitudinally at diagnosis, complete remission and relapse. Following thawing of frozen aspirates, the cells were quantified and immortalized Raji cells were added to the sample to achieve an approximate 1% spike in cell population. Known heterozygous SNVs within the Raji cells served as a positive control for cell type identification and a way to assess allele dropout in the workflow. Cell suspensions were then emulsified and barcoded with our workflow prior to bulk preparation of the final sequencing libraries. Total workflow time for each sample was less than two days. MiSeq runs generating 250 bp paired-end reads were performed for each of the three samples that were barcoded. On average, 74.7% of the reads (MAPQ > 30) were associated with a cell barcode and correctly mapped to one of the 62-targeted loci (Fig. 3a). Performance of the panel across the targeted loci is shown in Supplementary Figure 2. The Raji cell spike in detection rate across the three sample runs averaged 2.4% and the average allele dropout rate, calculated from two separate heterozygous *TP53* SNVs present in the Raji cells, was 7.0% (Fig. 3a). The allele dropout rate represents the percentage of cells within a run, averaged across the two loci, where the known heterozygous SNV was incorrectly genotyped as either homozygous wild type or homozygous mutant.

**Figure 3.**
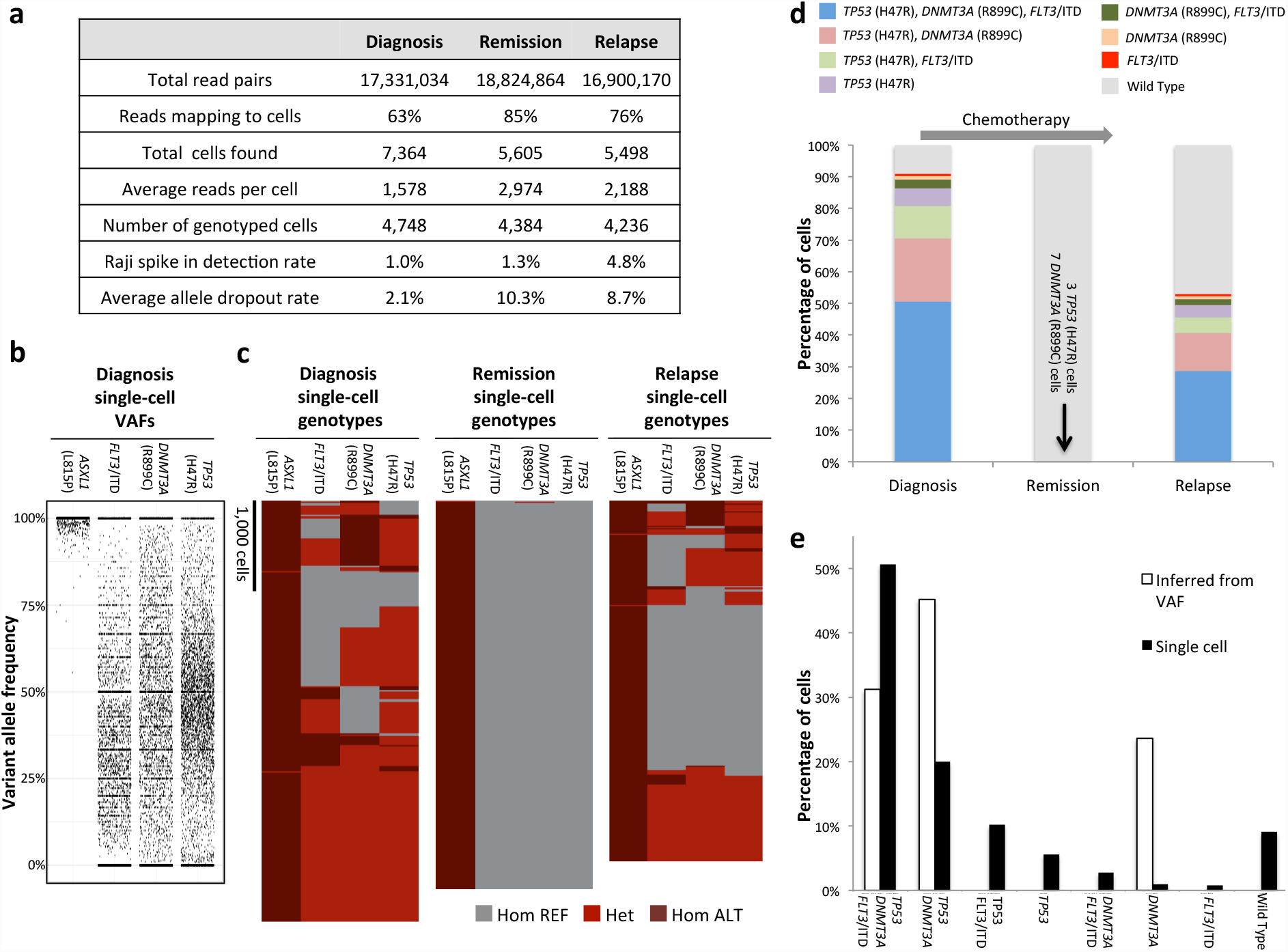
Analysis of AML clonal architecture. (a) Table displaying key metrics from the diagnosis, remission and relapse single-cell DNA sequencing runs from one patient. (b) Diagnosis sample single-cell VAFs for each of the 4 non-synonymous mutations identified for this patient. (c) Heat maps denoting single-cell genotypes for the three longitudinal patient samples. The presence of a heterozygous alternate (ALT) allele is shown in red. Homozygous alternate alleles are shown in dark red and reference alleles are depicted in gray. (d) Clonal cell populations identified from clinical bone marrow biopsies taken at the time of diagnosis, remission and relapse. Wild Type indicates cells that had reference genome sequence for *TP53*, *DNMAT3A* and *FLT3*, but were homozygous for the *ASXL1* (L815P) mutation. (e) Comparison of single-cell sequencing data from the diagnosis sample obtained from our workflow and a simple clonal inference of the diagnosis cell clonal populations produced from the bulk VAFs. Non-patient Raji cells have been removed for the analyses in (c), (d) and (e).

### Single-cell variant calling and clonal analysis of AML

Using standard genotype calling algorithms (See Materials and Methods), we identified a total of 17 variant alleles for this patient (Supplementary Table 2 and Supplementary Figure 3). While 13 of these variants occurred in noncoding DNA, three non-synonymous SNVs were found in coding regions of *TP53* (H47R), *DNMT3A* (R899C) and *ASXL1* (L815P) from all three longitudinal samples (Fig. 3c,d). *ASXL1* (L815P) is a previously reported common polymorphism (dbSNP: rs6058694) and was likely present in the germline since it was found in all cells throughout the course of the disease (*16*). Additionally, a 21 bp internal tandem duplication (ITD) in *FLT3* was detected in cells from the diagnosis and relapse samples. *FLT3*/ITD alleles are found in roughly a quarter of newly diagnosed adult AML patients and are associated with poor prognosis (*15, 17, 18*). A total of 13,368 cells (4,456 cells per run average) were successfully genotyped at the four variant genomic loci (Fig. 3a,b,c). A comparison of the clonal populations from the diagnosis, remission and relapse samples indicates that the patient initially achieved complete remission, although having 10 mutant cells demonstrates the presence of minimal residual disease (MRD) at this time point (Fig. 3d). Despite the initial positive response to therapy, the reemergence of the clones present at diagnosis in the relapse sample indicates that it was ineffective at eradicating all of the cancer cells and, in this instance, did not dramatically remodel the initial clonal architecture of the tumor. Single-cell sequencing of additional cells from the remission sample would likely be required to test this hypothesis and identify additional MRD clones.

To assess the performance of our single-cell approach relative to conventional next generation sequencing (see Materials and Methods), we obtained bulk variant allele frequencies (VAFs) for the relevant mutations in two of the biopsy samples. The bulk VAFs were comparable to the VAFs acquired from our single-cell sequencing workflow when the barcode identifiers are removed and the reads are analyzed in aggregate (Supplementary Figure 4). We next used the bulk sample VAFs to infer clonal architecture and compare it to the clonal populations obtained with our single-cell sequencing approach. The simplest model of inferred clonality predicts a significant *DNMT3A* (R899C) single mutant population indicative of founder mutation status (Fig. 3e). Interestingly, the single-cell sequencing data does not support this model as only a relatively small *DNMT3A* single mutant population is observed and this population is at a frequency that can be explained by allele dropout. In contrast, our results suggest that the SNV in *TP53* could be the founding mutation since the size of the *TP53* (H47R) single mutant clone is larger than what would be expected from allele dropout. Our single-cell approach also unambiguously identified the *TP53*, *DNMT3A* and *FLT3*/ITD triple mutant population as the most abundant neoplastic cell type in the diagnosis and relapse samples (Fig. 3d). Moreover, the identification of this clone strongly supports a model where the mutations were serially acquired during the progression of the disease.

### Clonal remodeling of an AML tumor at relapse

To further investigate the ability of high-throughput single-cell sequencing to accurately characterize the clonal architecture of tumors, we analyzed diagnosis and relapse bone marrow aspirate samples from a second patient with cytogenetically normal AML. Using our variant calling approach, 20 genetic variants were confidently identified and, of those, 5 were non-synonymous mutations (Supplementary Table S4). *ASXL1* (L815P) was again identified as a non-synonymous polymorphism in this patient, further validating its status as a common variant. We focused subsequent analysis on the disease relevant mutations that were also identified in bulk sequencing of the diagnosis sample, *IDH2* (R140Q), *NRAS* (G13R) and *ASXL1* (G646fs). A total of 2,850 cells were accurately genotyped at these three variant genomic loci (Fig. 4a). The Raji cell spike in detection rate for these runs averaged 1.5% and the average allele dropout rate, calculated from the Raji cells, was 8.5%.

**Figure 4.**
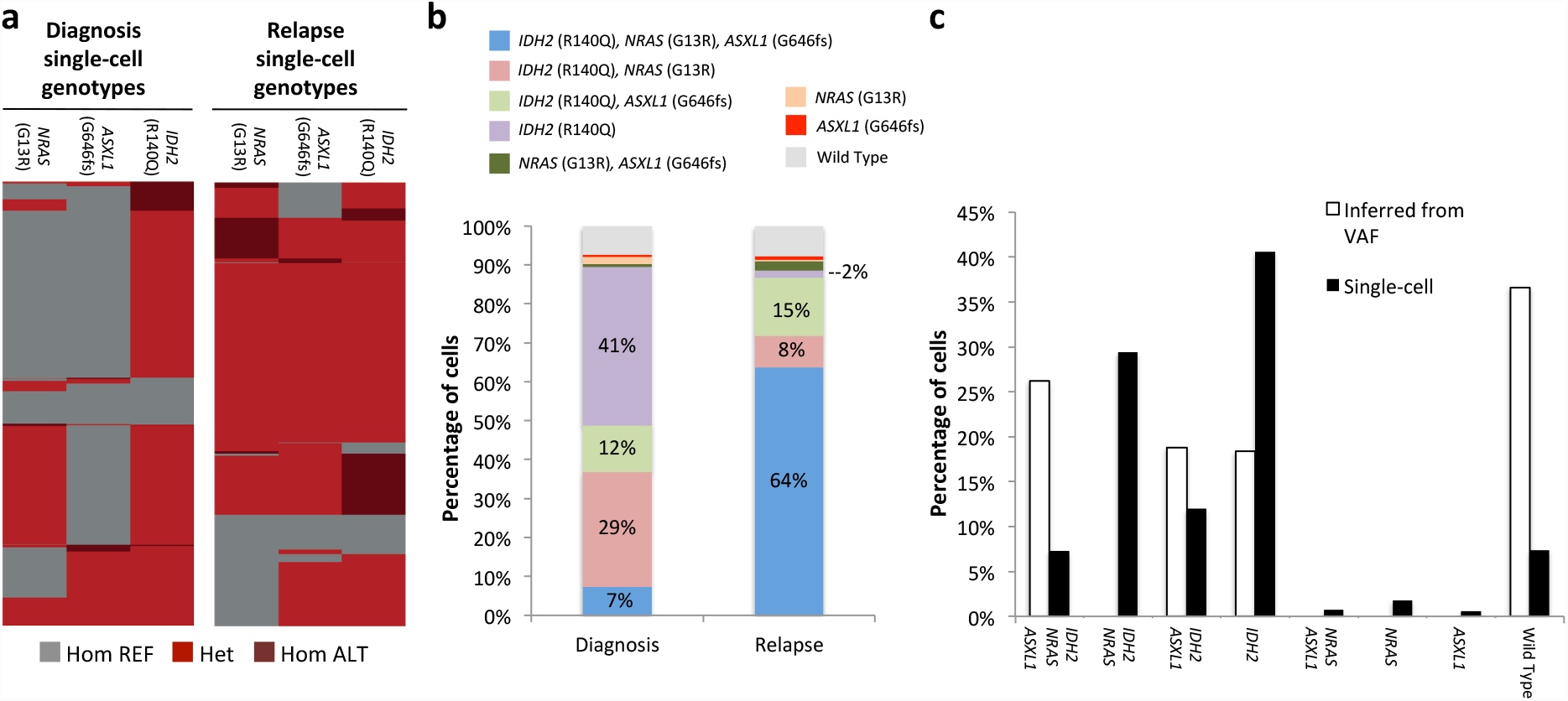
Clonal remodeling of an AML tumor. (a) Heat maps denoting single-cell genotypes for the diagnosis and relapse samples. The presence of a heterozygous alternate (ALT) allele is shown in red. Homozygous alternate alleles are shown in dark red and reference alleles are depicted in gray. (b) Clinical bone marrow biopsies taken at the time of diagnosis and relapse show substantial changes in clonal distribution with single-cell sequencing. Wild Type indicates cells that had reference genome sequence for *IDH2*, *ASXL1* and *NRAS* (c) Comparison of single-cell sequencing data from the diagnosis sample obtained from our workflow and a simple clonal inference of the diagnosis cell clonal populations produced from the bulk VAFs. Non-patient Raji cells have been removed from these datasets.

Using the single-cell genotype calls on individual cells, the clonal composition of both diagnosis and relapse tumors were reconstructed, as shown in Figure 4b. Strikingly, there were a number of clones that significantly expanded or contracted during the course of the disease. The most dramatic of these changes were an expansion of the triple mutant *IDH2* (R140Q) *NRAS* (G13R) *ASXL1* (G646fs) clone from 7% at diagnosis to 64% at relapse and a reduction in the *IDH2* (R140Q) single mutant population from 41% at diagnosis to 2% at relapse. These changes cannot be explained technical noise since the allele dropout rates were almost identical between the two sample runs (diagnosis = 8.0%, relapse = 9.0%). Additionally, if these clonal distribution changes were systematic and technical in nature, we would have expected to see similar changes in the first patient, where the diagnosis and relapse samples were almost identical in composition. One possible factor contributing to the clonal evolution seen in the second patient is the extended 3-year time period that elapsed between remission and relapse.

Lastly, we used VAFs generated from bulk sequencing to infer clonal architecture of the diagnosis sample. This inferred architecture differed substantially from the single-cell sequencing derived populations (Fig. 4c). Notably, the *IDH2* (R140Q) *NRAS* (G13R) clone was not predicted from the bulk sequencing VAFs, yet it represented 29% of the diagnosis tumor defined by single-cell DNA sequencing. The *IDH2* (R140Q) *NRAS* (G13R) population shrinks to 8% of the tumor at the relapse time point where the triple mutant *IDH2* (R140Q) *NRAS* (G13R) *ASXL1* (G646fs) clone expands to 64%. Consequently, it is not likely that the *IDH2* (R140Q) *NRAS* (G13R) clone is a result of *ASXL1* (G646fs) allele dropout in the triple mutant cells given the much greater size of the triple mutant cell population at relapse and similar observed allele dropout rates. This provides another clear example of the high-resolution clonal architecture uncovered by single-cell DNA analysis that is missed with bulk sequencing data.

## Conclusions

Our method enables rapid and cost-effective targeted genomic sequencing of thousands of AML cells in parallel – something that is not feasible with existing technologies. Applying this approach to the study of larger AML patient populations will likely lead to correlations between clonal heterogeneity and clinical outcomes. Although we focused on AML in this study, our method should be applicable to other cancer cell types and profiling of solid tumors that have been dissociated into single-cell suspensions. This capability is poised to complement an increased scientific appreciation of the role that genetic heterogeneity plays in the progression of many cancers as well as a desire by clinicians to make personalized medicine a widespread reality.

## MATERIALS & METHODS

### Cell and patient samples

All patient samples were collected under an IRB approved protocol and patients singed the consent for sample collection and analysis. The protocol adhered to the Declaration of Helsinki. The clinical AML samples presented in Figure 3 were obtained from a 66 year-old man that was diagnosed with AML, French-American-British (FAB) classification M5. Pre-treatment diagnostic bone marrow biopsy showed 80% myeloblast and cytogenetic analysis showed normal male karyotype. He received an induction chemotherapy consisted of fludarabine, cytarabine and idarubicin. Day 28 bone marrow aspiration showed morphological complete remission (CR). He received additional 2 cycles of consolidation therapy with the same combination but approximately 3 months after achieving CR, his AML relapsed with 48% blast. He was subsequently treated with azacitidine and sorafenib chemotherapy and achieved second CR. He then underwent allogeneic stem cell transplant from his matched sibling but approximately 2 months after transplant, his disease relapsed. He was subsequently treated with multiple salvage therapies but he passed away from leukemia progression approximately 2 years from his original diagnosis. Bone marrow from original diagnosis, first CR, and first relapse were analyzed.

The second patient analyzed and presented in Figure 4 was a 65 year-old man diagnosed with AML having myelodysplastic changes, 44% myeloblast, and cytogenetic analysis showed a normal karyotype. Bulk sequencing VAFs for *IDH2* (R140Q)*, NRAS* (G13R) and *ASXL1* (G646fs) were 31.7%, 13.1% and 22.5%, respectively in the diagnosis sample. The patient received induction chemotherapy with cladribine and cytarabine and achieved CR at day 28. He completed consolidation therapy for 18 cycles and then received maintenance therapy with decitabine for additional 12 cycles.

Approximately 3 years after achieved CR, patient relapsed with 20% blast. Tumor cell samples were not available for the remission time point of this patient.

Raji B-lymphocyte cells were cultured in complete media (RPMI 1640 with 10% fetal bovine serum (FBS), 100 U/ml penicillin, and 100 µg/ml streptomycin) at 37°C with 5% CO_2_. Cells were pelleted at 400 g for 4 min and washed once with HBSS and resuspended in PBS that was density matched with OptiPrep (Sigma-Aldrich) prior to encapsulation in microfluidic droplets. Frozen bone marrow aspirates were thawed at the time of cell encapsulation and resuspended in 5 ml of FBS on ice, followed by a single wash with PBS. All cell samples were quantified prior to encapsulation by combining 5 µl aliquots of cell suspension with an equal amount of trypan blue, then loaded on chamber slides and counted with the Countess Automated Cell Counter (ThermoFisher). The Raji cells were added to the bone marrow cell samples to achieve a ~1% final spike-in concentration.

### Fabrication and operation of microfluidic devices

We performed the microfluidic droplet handling on devices made from polydimethylsiloxane (PDMS) molds bonded to glass slides; the device channels were treated with Aquapel to make them hydrophobic. The PDMS molds were formed from silicon wafer masters with photolithographically patterned SU-8 (Microchem) on them. We operated the devices primarily with syringe pumps (NewEra), which drove cell suspensions, reagents and fluorinated oils (Novec 7500 and FC-40) with 2–5% PEG-PFPE block-copolymer surfactant into the devices through polyethylene tubing (*19*). Merger of the cell lysate containing droplets with the PCR reagent/barcode bead droplets was performed using a microfluidic electrode.

### Generation of barcode containing beads

Barcoded hydrogel beads were made as previously reported in Klein et al(*6*). Briefly, a monomeric acrylamide solution and an acrydite-modified oligonucleotide were emulsified on a dropmaker with oil containing TEMED. The TEMED initiates polymerization of the acrylamide resulting in highly uniform beads. The incorporated oligonucleotide was then used as a base on which different split-and-pool generated combinations of barcodes were sequentially added with isothermal extension. Targeted gene-specific primers were phosphorylated and ligated to the 5’ end of the hydrogel attached oligonucleotides. ExoI was used to digest non-ligated barcode oligonucleotides that could otherwise interfere with the PCR reactions. Because the acrydite oligo also has a photocleavable linker (required for droplet PCR), barcoded oligonucleotide generation could be measured. We were able to convert approximately 45% of the base acrydite oligonucleotide into full-length barcode with gene specific primers attached. Single bead sequencing of beads from individual bead lots was also performed to verify quality of this reagent.

### Cell encapsulation and droplet PCR

Following density matching, cell suspensions were loaded into 1 ml syringes and co-flowed with an equal volume of lysis buffer (100 mM Tris pH 8.0, 0.5% IGEPAL, proteinase K 1.0 mg/ml) to prevent premature lysing of cells(*20*). The resultant emulsions were then incubated at 37°C for 16–20 hours prior to heat inactivation of the protease.

Droplet PCR reactions consisted of 1X Platinum Multiplex PCR Master Mix (ThermoFisher), supplemented with 0.2 mg/ml RNAse A. Prior to thermocycling, the PCR emulsions containing the barcode carrying hydrogel beads were exposed to UV light for 8 min to release the oligonucleotides. Droplet PCR reactions were thermocycled with the following conditions: 95°C for 10 min, 25 cycles of 95°C for 30 s, 72°C for 10 s, 60°C for 4 min, 72°C for 30 s and a final step of 72°C for 2 min. Single-cell TaqMan reactions targeting the SRY locus were performed as previously described(*12*).

### DNA recovery and sequencing library preparation

Following thermocycling, emulsions were broken using perfluoro-1-octanol and the aqueous fraction was diluted in water. The aqueous fraction was then collected and centrifuged prior to DNA purification using 0.63X of SPRI beads (Beckman Coulter). Sample indexes and Illumina adaptor sequences were then added via a 10 cycle PCR reaction with 1X Phusion High-Fidelity PCR Master Mix. A second 0.63X SPRI purification was then performed on the completed PCR reactions and samples were eluted in 10 μl of water. Libraries were analyzed on a DNA 1000 assay chip with a Bioanalyzer (Agilent Technologies), and sequenced on an Illumina MiSeq with either 150 bp or 250 bp paired end multiplexed runs. A single sequencing run was performed for each barcoded single-cell library prepared with our microfluidic workflow. A 5% ratio of PhiX DNA was used in the sequencing runs.

### Analysis of next generation sequencing data

Sequenced reads were trimmed for adapter sequences (cutadapt(*21, 22*)), and aligned to the hg19 human genome using bwa-mem(*23, 24*) after extracting barcode information. After mapping, on target sequences were selected using standard bioinformatics tools (samtools(*25*)), and barcode sequences were error corrected based on a white list of known sequences. The number of cells present in each tube was determined based on curve fitting a plot of number of reads assigned to each barcode vs. barcodes ranked in decreasing order, similar to what described in Macosko et al. (*8*). The total number of cells identified in this manner for a given sample run are presented in Figure 3a as “Total cells found”. A subset of these cells was then identified that had sufficient sequence coverage depth to call genotypes at the non-synonymous variant positions identified in *TP53, ASXL1, FLT3* and *DNMT3A*. This subset of cells is presented as “Number of genotyped cells” in Figure 3a.

GATK 3.7(*26*) was used to genotype diagnosis samples with a joint-calling approach. The quality score of known Raji cell mutations was used to set a minimum threshold for variant calling in patient cells. For patient 1 (Fig. 3), the presence of these variants as well as the potential *FLT3*/ITD were called at a single cell level across the three samples using Freebayes(*27*). Genotype cluster analysis was performed using heatmap3 for R(*28*). The non-patient Raji cell spike in populations were removed for these analyses.

### Bulk sequencing using capture targeted sequencing

We designed a SureSelect custom panel of 295 genes (Agilent Technologies, Santa Clara, CA) that are recurrently mutated in hematologic malignancies (Supplementary Table 3). Extracted genomic DNA from bone marrow aspirates was fragmented and bait-captured according to manufacturer protocols. Captured DNA libraries were then sequenced using a HiSeq 2000 sequencer (Illumina, San Diego, CA) with 76 bp paired-end reads.

## Acknowledgements

We thank Bill Hyun and Hannah Viernes for comments on the manuscript.

## Author Contributions

D.J.E. and M.P. conceived the technology, designed the study/experiments, analyzed the data and wrote the manuscript. D.J.E. performed the AML sample runs and prepared the libraries for sequencing. M.P. developed the bioinformatics analysis pipeline and performed bioinformatics analysis on next-generation sequencing data. A.S. designed and fabricated the microfluidic devices. S.T. and W.O. generated the barcoded beads. S.T., K.G., J.J., T.C. and K.W.J. optimized droplet biochemistry and developed the targeted sequencing panel. R.D. assisted with bioinformatics analysis on next generation sequencing data. K.T., J.M. and H.K. provided AML research samples and assisted with the interpretation of the sequencing data. K.T. and P.A.F. performed bulk sequencing of the same AML samples. All authors read and approved the final manuscript.

## Supplementary Information

Table S1. *AML targeted sequencing panel.* List of the 25 genes and genomic locations sequenced with the 62 amplicon targeted sequencing panel.

Figure S1. *Uniformity of amplification across the 62 amplicon AML panel.* The percentage of total reads per cell is plotted for each of the 62 amplicons in the targeted AML panel. Each point in the plot represents the data from a single-cell. Blue colored data points were generated for droplet amplification conditions prior to uniformity optimization. Red data points were generated following optimization of thermocycling conditions and reaction components. For each of the two separate conditions the same 250 cells are shown across all of the different sequenced loci.

Figure S2. *Single-cell targeted panel amplification performance.* Histograms showing the percentage of the 62 target amplicons present at either 10X (dark gray) or 20X (light gray) coverage across the cells identified in our bioinformatics pipeline. Data is shown for single-cell sequencing of the diagnosis, remission and relapse samples.

Table S2. *List of the sequence variants identified from the patient 1 AML diagnosis sample.* From this list, four non-synonymous coding region variants were identified in *ASXL1*, *TP53*, *DNMT3A* and *FLT3*.

Figure S3. *Single-cell variant allele frequencies.* Single cell VAFs are plotted for all 17 of the identified genetic variants found in the diagnosis, complete remission and relapse samples.

Figure S4. *Comparison of bulk and pseudo bulk VAFs.* Variant allele frequencies for mutations in *TP53*, *DNMT3A* and *FLT3* are displayed for diagnosis, remission and relapse samples. VAFs obtained from traditional bulk next generation sequencing are displayed as open bars. The black bars represent the VAFs obtained from our single-cell sequencing workflow; however, the barcode identifiers have been removed and the reads have been analyzed as a single population to give a “pseudo-bulk” frequency. Bulk sequencing was not performed on the remission sample; therefore, VAF data is not available for this timepoint.

Table S3. *List of the 295 genes that were targeted for bulk sequencing.*

Table S4. *List of the sequence variants identified from the patient 2 AML diagnosis sample.* From this list, five non-synonymous coding region variants were identified in *ASXL1*, *IDH2*, *NRAS* and *SRSF2*. These variants were identified with a targeted single-cell sequencing panel that was slightly modified from the panel shown in Table S1 with amplicons covering *CBL*, *MIR142*, *TET2*, *PHF6*, *RAD21* and *STAG2* removed. The *SRSF2* mutation was not considered for further analysis due to the majority of cells identified not having any sequence coverage at this position.

